# Active Sinking Particles: Sessile Suspension Feeders significantly alter the Flow and Transport to Sinking Aggregates

**DOI:** 10.1101/2021.08.05.455190

**Authors:** Deepak Krishnamurthy, Rachel Pepper, Manu Prakash

## Abstract

Sinking or sedimentation of biological aggregates plays a critical role in carbon sequestration in the ocean and in vertical material fluxes in waste-water treatment plants. In both these contexts, the sinking aggregates are “active,” since they are biological hot-spots and are densely colonized by microorganisms including bacteria and sessile protists, some of which generate feeding currents. However, the effect of these feeding currents on the sinking rates, trajectories, and mass transfer to these “active sinking particles,” has not previously been studied. Here we use a novel scale-free vertical-tracking microscope (a.k.a. Gravity Machine, Krishnamurthy et al. “Scale-free vertical tracking microscopy.” Nature Methods (2020)) to follow model sinking aggregates (agar spheres) with attached protists (*Vorticella convallaria*), sinking over long distances while simultaneously measuring local flows. We find that activity due to attached *Vorticella* cause substantial changes to the flow around aggregates in a dynamic manner and reshape mass transport boundary layers. Further, we find that activity-mediated local flows along with sinking significantly changes how aggregates interact with the water-column at larger scales by modifying the encounter and plume cross-sections and by inducing sustained aggregate rotations. In this way our work suggests an important role of biological activity in understanding the growth, degradation, composition and sinking speeds of aggregates with consequences for predicting vertical material fluxes in marine, freshwater and man-made environments.

**Significance Statement:** Sinking aggregates are a critical part of aquatic ecosystems. Plentiful sinking aggregates account for the majority of carbon sequestration in the oceans. These aggregates are densely colonized by microorganisms, including many that generate feeding currents. Utilizing a novel instrument for high resolution imaging of sinking particles, we demonstrate that these feeding currents significantly change how water flows near the aggregates. We show that these changes in flow are likely to affect aquatic system processes, including aggregation rates, degradation rates, sinking speeds, and aggregate composition. Our work provides a starting point for exploring the larger-scale implications of attached organisms on these system processes, which, in turn, are critical for understanding carbon sequestration in the oceans or efficiency in waste-water treatment plants.

## 2 Introduction

Suspended aggregates are key parts of natural and man-made aquatic ecosystems. In particular, macroaggregates (> 500 *µ*m), known as marine snow, lake snow, and river snow, can dominate nutrient, carbon, and element cycling in aquatic environments [1, 2, 3, 4, 5, 6, 7]. In the ocean marine snow aggregates are primarily responsible for vertical material fluxes from the surface mixed layer to the deep ocean, thus playing a critical role in marine carbon sequestration [5, 1, 6]. Similar aggregates are also a key part of activated sludge wastewater treatment facilities, where the sinking out of flocs is one of the main methods for removing organic debris and contaminants from the water [8]. Understanding aggregation rates, degradation rates, sinking speeds, and composition of aggregates in aquatic environments is critical for understanding vertical fluxes of carbon and nutrients in these diverse ecosystems [7, 1, 9]. These rates can in-turn depend on several biotic and abiotic factors. The local hydrodynamic environment is one such factor: encounter rates, which determine particle size distributions and sinking rates, can depend sensitively on fluid flow near the aggregate, e.g., local shear, or the flow regime (Reynolds number). [9, 10].

In marine, freshwater, and waste-water contexts suspended aggregates are biological hotspots: they are highly enriched in carbon and other nutrients, compared to the surrounding water, and are, therefore, an important micro-habitat for aquatic organisms [11, 12, 13, 7, 2, 14] (Fig. 1A, B, C). Marine and freshwater aggregates are typically densely colonized by bacteria, flagellates, ciliates, and are consumed by larger zooplankton [11, 15, 13, 7, 2]. Similarly, wastewater flocs are also densely colonized by bacteria and protists [16]. Sessile ciliates attached to flocs are especially important for effective wastewater treatment, and the particular species composition is used as an indicator of the stage and health of the effluent [17, 16].

**Figure 1:**
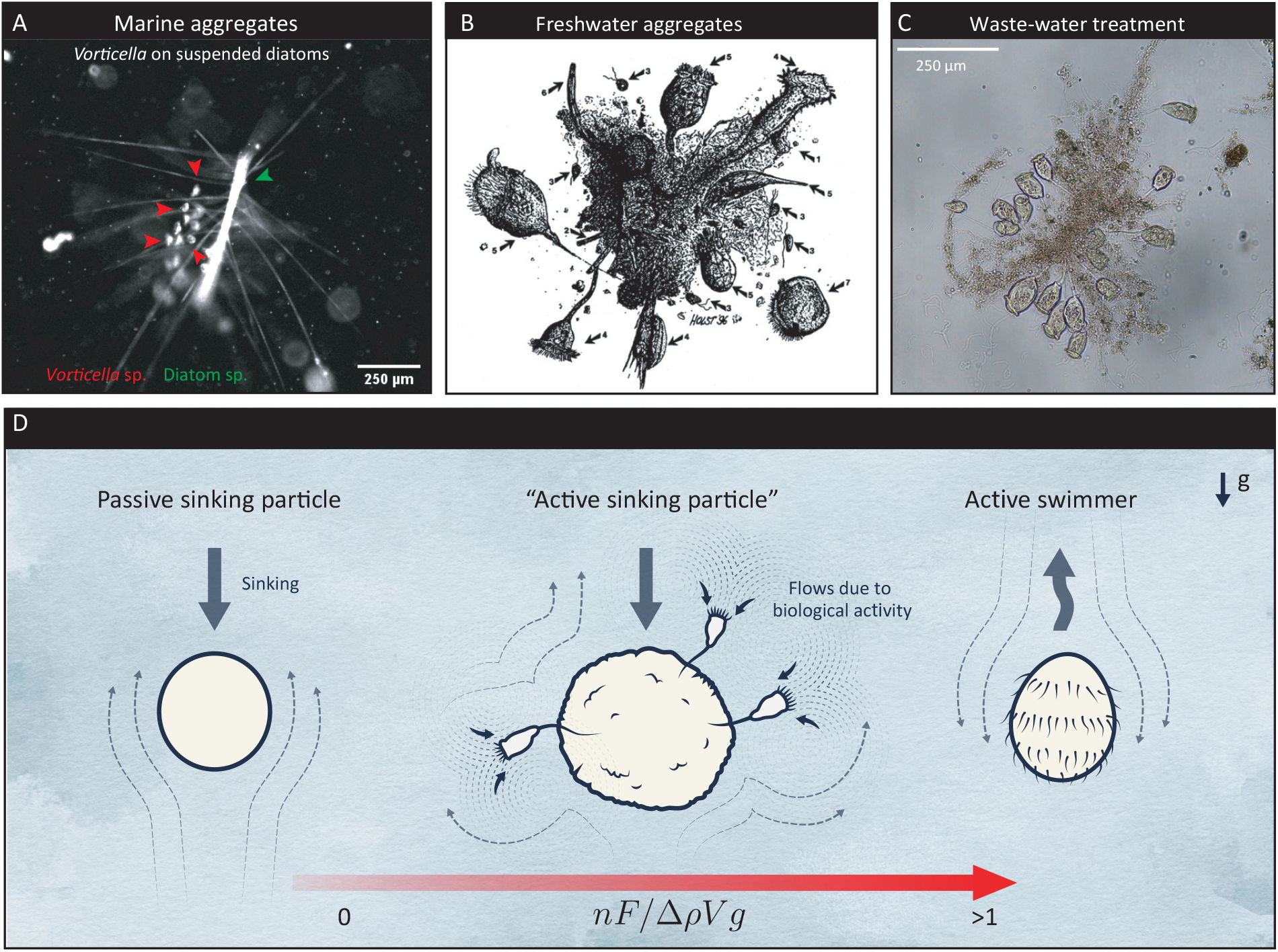
“Active sinking particles” in natural and man made ecosystems. **(A)** A snapshot of a natural marine aggregate including a diatom chain (red arrows) colonized by suspension feeders (green arrows). **(B)** A freshwater aggregate colonized by various suspension feeders including *Vorticella* species, *Stentor coeruleus*, and flagellates, such as *Bodo* species [15]. Image from Zimmerman-Timm *et al*. [15]. **(C)** An aggregate from the Tacoma Central waste-water treatment plant showing dense colonization by suspension feeding microorganisms. **(D)** The examples in (A), (B) and (C) constitute “active sinking particles” wherein the aggregate’s flow and mass transport characteristics have contributions both due to sinking as well as the active flows created by sessile microorganisms. The ratio of the effects of activity and sinking can be parametrized by the dimensionless ratio: 𝒜 ∼ *nF/*Δ*ρV g*, where *n* is the no:of active entities (*Vorticella* in our study), *F* is the force exerted on the fluid due to each active entity, Δ*ρ* is the density mismatch between the aggregate and fluid, *V* the aggregate volume and *g* the acceleration due to gravity.

How does biological activity of a sessile attached feeder fundamentally alter the hydrodynamic environment surrounding these sinking aggregates? Many of the organisms found on aggregates, including protists and nanoflagellates, generate a feeding current to draw in food from the surrounding fluid [15, 18, 13, 7]. These feeding currents can have velocities on the same order as near-field flows due to sinking (typical feeding current velocities 0.1–1*mms*^*−*1^ [19, 20] and sinking speeds of aggregates 𝒪 (1 mms^−1^)) [18]. We term these aggregates as *“active sinking particles,”* with the local flow fields determined both by sinking and biologically-generated flows. It is important to understand how this combination of sinking and biological activity affects mass transport, encounter rates with other aggregates, colonization of bacteria and other suspension feeders, as well as growth, degradation and sinking rates, yet these questions have not been systematically investigated so far. We have found only only one study of this potentially-important phenomenon; the feeding currents of attached nanoflagellates was shown to dramatically increase aggregation rates for small (< 20 *µ*m) particles, but direct measurement of flows are needed to confirm the mechanism and to extend to both larger organisms and larger aggregates [18].

The sinking and mass transport to “active sinking particles” in their ecological context is a complex, multi-scaled process. For instance, marine snow particles of size-scale 𝒪 (1 mm) sink at rates of 𝒪 (100 m*/*day) while undergoing myriad transformations due to both physical and biological influences [21, 22, 5]. *In situ* measurements of such processes are naturally challenging, and earlier efforts in the laboratory have used techniques including tethering aggregates and using a fixed background flow to simulate freely sinking conditions [6, 23]. However, practical constraints posed by these experimental setups have made it challenging to bring modern microscopy techniques into the realm of this problem. Direct, real-time observations of near-field flows (single cell to aggregate scale), as well as dynamics of freely sinking aggregates, over both short and long time-scales, are currently lacking.

In this work, we present a novel experimental system and measurements combining agar aggregates colonized by *Vorticella convallaria* (our model “active sinking particles”), and Scale-free Vertical Tracking Microscopy, aka Gravity Machine [24]. This recent advance in tracking microscopy allows freely-sinking particles to be automatically tracked using a circular “hydrodynamic treadmill.” Using this novel method we present the first detailed measurements and characterizations of near-field flows around “active sinking particles” and find that they can be significantly modified by the presence of one or more *Vorticella* cells. We also find that *Vorticella* modify aggregate dynamics by causing sustained rotations, with implications for mass transport and aggregate transformations over long times. Finally, we show how the presence of one or more *Vorticella* can affect aspects of mass transport to the aggregate including the encounter region and plume cross-sections, important parameters in estimating mass transport and aggregate growth and degradation rates. Overall, our work introduces a new experimental method allowing novel multi-scale measurements of “active sinking particles,” and highlights the importance of biological activity in shaping the nearfield flows and mass transport to sinking aggregates.

## 3 Results

### 3.1 Conceptual framework

Active sinking particles lie on the continuum spanned, on the one hand, by passive sinking particles that have no biological activity, and, on the other, by active swimmers under the influence of gravity (Fig. 1). This continuum can be parameterized by a non-dimensional ratio of stresses due to activity and buoyancy. The active stress can be written as 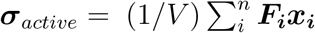 [25], where *n* is the number of active entities (*Vorticella* in our case) whose influence of fluid is approximated as point-forces (Stokeslets) of strength ***F*** _*i*_ [26], ***x***_*i*_ are their distances from the aggregate center, and *V* is the aggregate volume. The stress due to gravity is Δ*ρV g/a*^2^ where Δ*ρ* is the density mismatch between the particle and ambient fluid, *g* is the acceleration due to gravity, and *a* is the aggregate radius. Thus, the dimensionless ratio of this active stress to the stress on the fluid due to buoyant forces then scales as: 𝒜 ∼ *nF/*Δ*ρV g*. Using this parametrization, passive sinking particles have 𝒜 = 0, active swimmers are typically characterized by 𝒜 ≳ 1, while active sinking particles have 0 < 𝒜 ≲ 1 (Fig. 1D).

### 3.2 Experimental system

To study the multi-scale process of microscale near-field flows and macroscale sinking dynamics we leveraged Scale-free Vertical Tracking Microscopy (SVTM) [24] that uses controlled rotation of a circular “hydrodynamic treadmill” to automatically keep small objects centered in the microscope field-of-view while allowing free movement with no bounds along the axis of gravity (Fig. 2A). SVTM allowed us to concurrently measure aggregate trajectories over meters (Fig. 2B), and capture images through video-microscopy at rates of 100 frames/s, (Fig. 2C, Movie S1). The optical resolution of the imaging system was ≈ 1*µm*, thus enabling resolution of flows and dynamics at the scale of single-cells colonizing the aggregate (Fig. 2 D). We observed sinking spherical millimeter-scale aggregates that were either bare or had one to several attached *V. convallaria*, a common species of sessile ciliates (see Materials and Methods for further details). *Vorticella* species are found abundantly on sinking aggregates in marine, freshwater, and man-made ecosystems [15, 16]. We then measured flow around active sinking particles using Particle Image Velocimetry (PIV) and particle pathline traces on short video segments (see Materials and Methods for further details).

**Figure 2:**
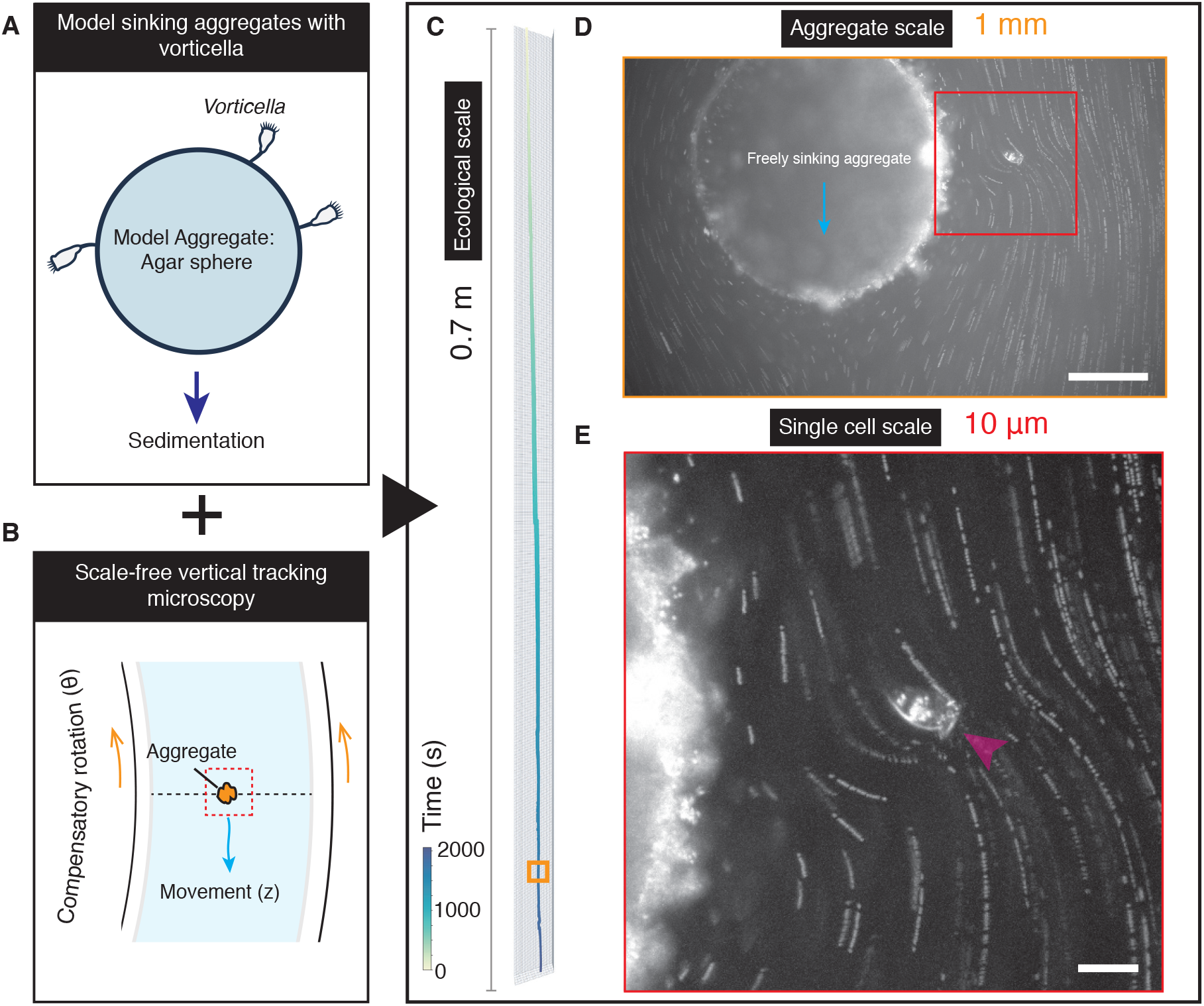
Scale-free Vertical Tracking Microscopy of freely sinking model aggregates with attached *Vorticella*. **(A)** Scale-free Tracking Microscopy using a “hydrodynamic treadmill” allows multi-scale tracking of aggregates: observations are made at microscale resolution while concurrently allowing free sinking and tracking over macro-scales. **(B)** Model sinking aggregates comprising of agar spheres of radius 0.5 *mm*, and sedimentation speed of ≈ 1 *mms*^−1^, colonized by *V. convalaria*. **(C)** A typical 3D trajectory of an aggregate showing sinking over 0.7 meters in 33 mins. In general this multi-scale tracking method results in tracks where the spatio-temporal resolution is microns and millseconds while the track extends over meters and hours. **(D)** Concurrent microscopy images of the aggregate captured in real-time at a rate of 30 Hz showing the aggregate and sessile suspension feeder. The snapshot corresponds to the orange box shown in (C). **(E)** Close-up view of the *Vorticella* cell demonstrating the fine optical resolution of the system. Arrow indicates the orientation of the cell and equivalently the direction in which stress is applied to the fluid.

### 3.3 *Vorticella* modify near-field flow and reshape boundary-layers

Attached *Vorticella* substantially changed the flows near sinking aggregates in ways that were variable and depended sensitively on the location and orientation of the attached organisms (representative examples in Fig. 3; all measured flow fields in Supplementary Fig. 1; N=12). A typical bare aggregate had relatively smooth flow (Fig. 3A,B) and both perpendicular and parallel flow near the particle that was similar the predictions of flow at zero Reynolds number around a sphere of the same size (Fig. 3C, Movie S2). As expected, tangential flow was fastest at the equator, and radial flow was fastest at the poles (Fig. 3C).

**Figure 3:**
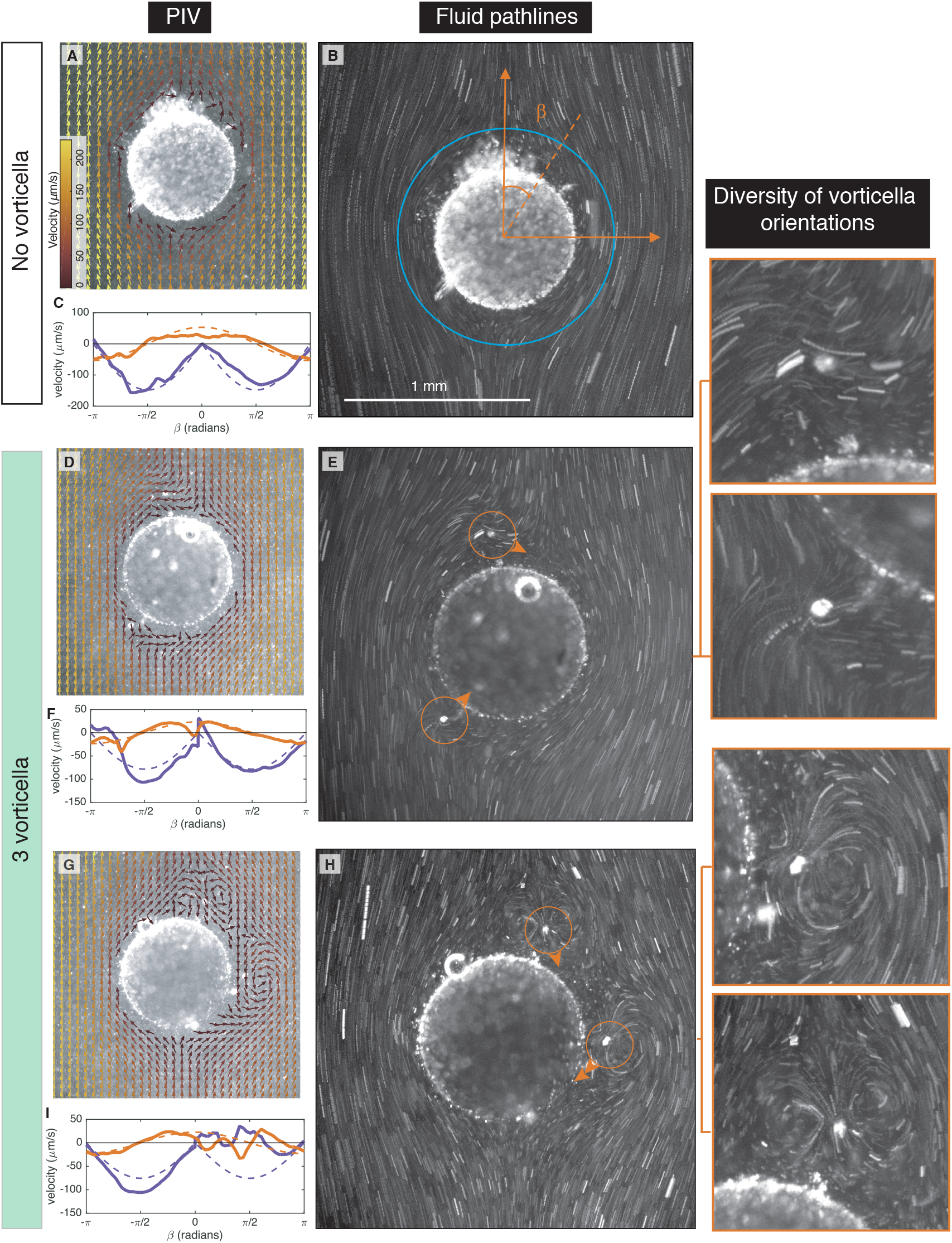
Flow around active sinking aggregates. Each row has measurements for a different sinking aggregate. **(A**,**D**,**G)** Measured flow velocities. Arrows indicated flow direction and color indicates speed. **(B**,**E**,**H)** Streaklines formed from tracer particles over ≈ 1 *s*. Circles indicate location of *Vorticella* and arrows indicated the direction of *Vorticella* forcing. **(C**,**F**,**I)** Measured velocities (solid lines) along a circle 200 *µ*m from the sphere surface (e.g. blue line in panel (B)). *β* is the polar angle indicated in panel (B), and is positive to the right of the sphere and negative to the left. Perpendicular and parallel components of the velocity are relative to the sphere surface. Dashed lines are calculated velocities for Stokes flow (zero Reynolds number) around a sphere. **Aggregate 1; top row; A-C**: No attached *Vorticella*; *a* = 380 *µ*m; *U* = 340 *µ*m*/*s; Re = 0.3. **Aggregate 2; middle row; D-F**: Three attached *Vorticella* (one out of view); *a* = 490 *µ*m; *U* = 200 *µ*m*/*s; Re = 0.2. **Aggregate 3; bottom row; G-I**: Three attached *Vorticella*; *a* = 490 *µ*m; *U* = 210 *µ*m*/*s; Re = 0.2. Representative snapshots of other experimentally-measured flow fields are shown in Supplementary Fig. 1.

On the other hand, Fig. 3 also shows two aggregates, each with three *Vorticella* attached, where the flow was modified by the *Vorticella*. In Fig. 3D,E,F, the modifications were somewhat subtle: streamlines were pulled toward the *Vorticella*, disrupting the background flow (Fig. 3D,E, Movie S3). The *Vorticella* in the bottom left, which was pulling fluid towards the particle, increased the radial flow, and caused the tangential flow to be reduced on one side of the organism and enhanced on the other side (Fig. 3F). The *Vorticella* near the north pole changed the local flow from primarily radial to primarily tangential (Fig. 3F). Since the *Vorticella* altered the flow velocities and gradients near the sphere, they caused the boundary layer to be thicker in some regions of the sphere (where velocities were reduced), and to be thinner in others (where velocities were increased). These changes were more substantial and wide spread than smaller scale changes induced by irregularities on the surface of a bare aggregate (Fig. 3C). In Fig. 3G,H,I, the modifications to the flow by the *Vorticella* were more dramatic: *Vorticella* on this aggregate caused large-scale eddies, which changed the flow structure, completely altering tangential and radial flow velocities near the sphere (Fig. 3G,H,I, Movie S3). In some regions, the flow direction was even reversed from what it would have been for a bare sinking aggregate (Fig. 3I). Again, the boundary layer near this sphere was substantially re-shaped in ways that would likely change both total mass transport to the sphere and which regions of the sphere have the highest rates of mass transfer. Such diversity of flow structures caused by *Vorticella* feeding flow is similar to that observed and predicted previously for *Vorticella*, and similar organisms, attached to surfaces and in ambient flow [27, 20, 28, 26].

To understand the relative effects of activity and sinking on mass transport from the aggregates, we can compare their relative contributions to the velocity within the mass transport boundary-layer. Advective effects of the flow dominate diffusive effects for the range of aggregate sizes, sinking speeds and even for transport of small molecules such as oxygen from the aggregate (Materials and Methods). This implies that the non-dimensional Péclet number (Pe = *Ua/D*), which quantifies the relative importance of these effects, is much greater than 1 (Materials and Methods). Here, *U* is velocity scale of the flow, *a* the aggregate radius and *D* the diffusivity of the molecule of interest. At such high Péclet number, the mass transport boundary layer is a thin region near the aggregate whose thickness is ∼ *a*Pe^−1*/*3^ [29]. The hydrodynamic effect of *Vorticella* on the fluid can be modelled as a point-force of magnitude *F* at a height *h* above the boundary [26]. The velocity scale due to *Vorticella* within this boundary layer is then *F/µh*. On the other hand, the velocity scale within the boundary layer due to sinking is 𝒪 (*U* Pe^−1*/*3^). Comparing these two scales in the vicinity of a *Vorticella* cell we obtain a critical sinking speed that separates the “activity-controlled” and “sinking-controlled” mass transport regimes:

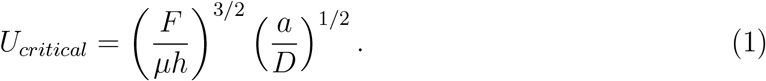

For sinking speeds above this scale, sinking is expected to dominate mass transport. Note that this picture is local, and the overall effect of the mass transport will depend on the number of *Vorticella*, as well as their orientation relative to the sinking direction of the aggregate. For instance, when *Vorticella* are oriented parallel to the local flow they can cause local flow reversals that completely disrupt the boundary layer structure and cause it to lift-off the aggregate surface (see for eg. Fig. 3G,H). Based on the measured parameters for *Vorticella* (*F* = 200 *pN, h* ∼ 100 *µm*, [28]), aggregate size *a* = 0.5 *mm*, and for transport of small-molecules like oxygen (*D* = 10^−9^ *m*^2^ *s*^−1^), we obtain *U*_*critical*_ = 63 *mm s*^−1^. This number is much higher than the sinking speeds in our experiments (which in-turn are representative of sinking speeds of real aggregates), indicating that activity dominates mass transport locally.

### 3.4 *Vorticella* modify encounter region and plume width

The above changes in near-field flows and mass transport of sinking aggregates due to attached organisms will have implications for many important processes, such as aggregate coagulation and growth rates, resulting size and shape of aggregates, aggregate sedimentation rates, the nutrient environment for organisms living on the aggregate, the plume left behind the aggregate as it sinks, and thus, chemotaxis of organisms to the aggregate [9, 10, 1, 30, 31, 32, 33, 7]. A full understanding of changes to mass transport requires full 3-dimensional flow fields and a numerical solution of the advection-diffusion equation (e.g., [34]), which are beyond the scope of this investigation. However, our experimental flow measurements are a key missing first piece in this analysis [35]. These flow fields also allow us to determine directly some of the changes to aggregate encounter rates and to the plume left behind by the aggregate.

We find that attached *Vorticella* significantly modify both the region of encounter below sinking aggregates and the shape of the plume left behind (Fig. 4). The encounter region below a sinking aggregate is the volume of fluid where particles will, eventually, come in to contact with the aggregate as it falls through the water column; the size and shape of this region is important for determining particle aggregation rates [10]. The plume is a volume of higher solute concentration left behind the aggregate as it sinks; bacteria and zooplankton may use the plume to find and colonize (or consume) sinking aggregates [6, 7, 33, 36, 37]. We estimated the width of the plume left behind by the aggregate as well as the width of the encounter region below the aggregate by following streamlines in our measured flow field; we also calculated this region analytically for Stokes and Oseen flow (more details in Materials and Methods). Stokes flow is accurate for zero Reynolds number, while Oseen flow makes adjustments for Reynolds numbers close to one [38]. Our example aggregates show that bare spheres have encounter regions and plumes fairly close to those predicted by Stokes and Oseen flow (Fig. 4A), while those with *Vorticella* have plumes and encounter regions with significant asymmetry and that can be both narrower and wider than those of bare spheres (Fig. 4B-E). Changes are particularly dramatic when the *Vorticella* cause recirculation in the flow (Fig. 4C-E). When extrapolated to distances far from the sphere (more details in Materials and Methods), we find that the width of the region encountered by the sphere and the width of the plume left behind are significantly changed by attached organisms, and that these changes lead to both thicker and thinner plumes and wider and narrower encounter regions (Fig. 4F,G). Each of our measurements are at a single time point and measure a 2D cross-section of a 3D flow. It is, therefore, possible that, for instance, in some cases that attached *Vorticella* cause the plume to be thinner in one dimension while thicker in another. Also, these changes are likely to be dynamic in time, leading to, e.g., a sometimes wider encounter region and sometimes narrower as time passes and *Vorticella* orientations and positions on the aggregate change.

**Figure 4:**
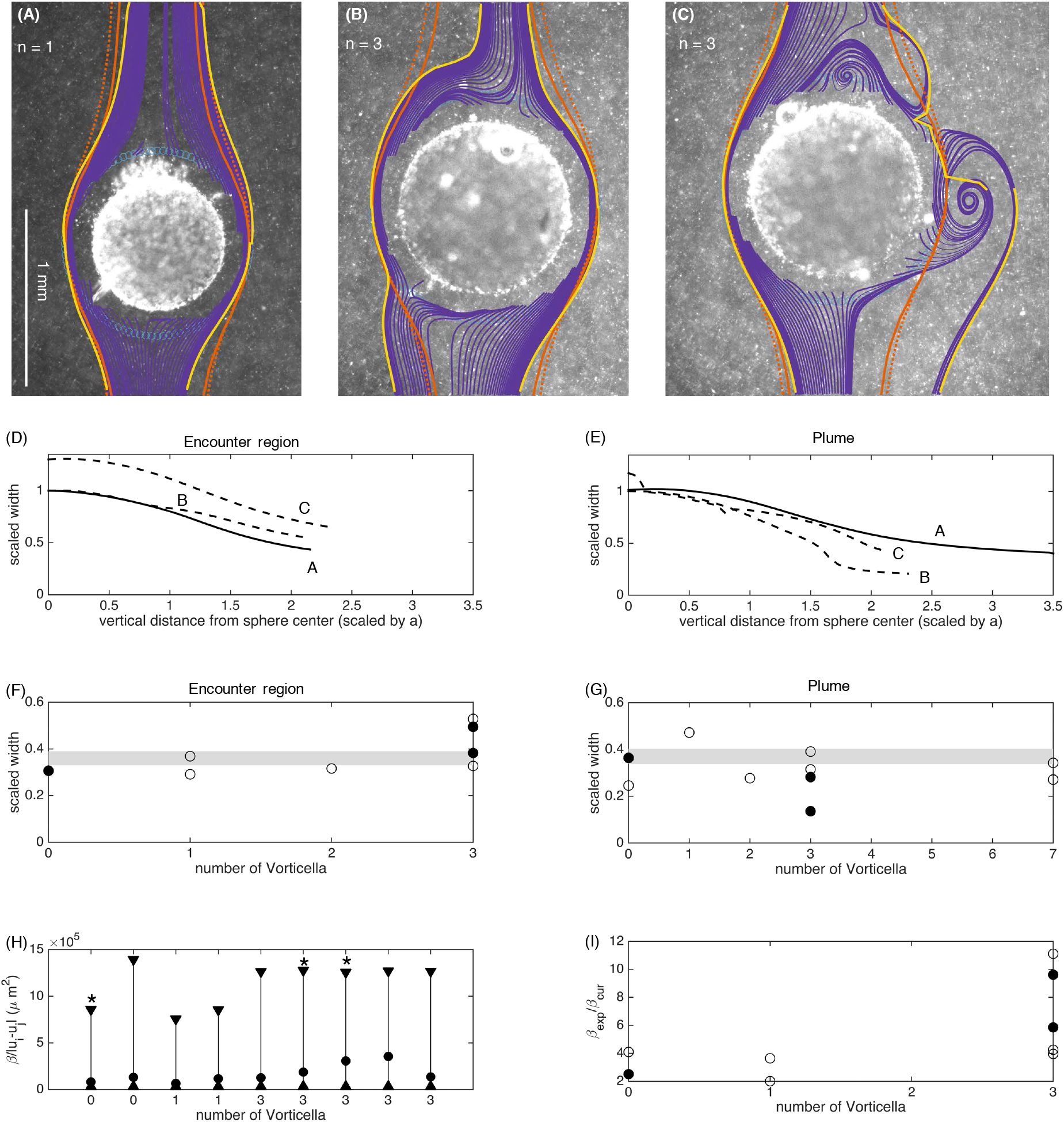
Encounter region and plume of sinking aggregates. **(A-C)** Streamlines around sinking aggregates (purple) calculated using experimentally-measured flow fields. Streamlines begin in a ring around the aggregate (blue circles) and are followed backward in time to find the encounter region for the aggregate (below the sphere) and forward in time to find the plume (above the sphere). Yellow lines indicates outer limits of these streamlines (e.g. the width of the plume and encounter region). Orange lines indicate theoretically calculated plume and encounter region widths, which were calculated for a sphere with matching radius and sinking speed. Of these, the solid lines indicate Stokes flow and dashed lines indicate Oseen flow. **(D**,**E)** Width of the intersection cross section and plume for the aggregates in panel (A) - (C). Letters indicate which panel matches which line and the solid line indicates that no *Vorticella* are attached. Widths are scaled by initial width at the equator of the sphere. **(F**,**G)** Long-distance encounter cross section and plume width (N = 9 for encounter region and N = 10 for plume). This width is calculated by continuing the experimentally-measured widths to *y* = ±20*a*. Solid circles indicate aggregates shown in panels (A) - (C) and shaded region indicates the range predicted by Oseen flow. **(H**,**I)** Particle coagulation kernels scaled by encounter velocity (N = 9). (H): The rectangular (upper triangles), experimental (circles), and curvilinear (lower triangles) kernels are compared for the parameters of each experimental aggregate. Stars indicate aggregates shown in panels (A) - (C). (I): The experimental coagulation kernel, *β*_*exp*_, divided by the curvilinear kernel, *β*_*cur*_ (N = 9). This indicates by how much encounter rates would be increased in our measured flow versus using the assumptions of Stokes flow. Solid circles indicate aggregates shown in panels (A) - (C).

We also used our long-distance encounter-width measurements (Fig. 4G) to predict how attached organisms change particle coagulation kernels. Particle encounter dynamics are included in aggregation models through these coagulation kernel that expresses encounter rate (in volume/time) [9]. The two most commonly used kernels are the rectilinear kernel:

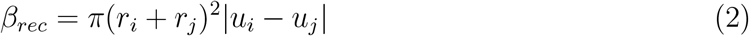

and the curvilinear kernel:

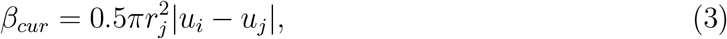

where *r*_*i*_ is the radius of the larger particle (here the sinking agar sphere), *r*_*j*_ is the radius of the smaller particle (here assumed to be 140 *µm* for simplicity - see Materials and Methods for further details), and *u*_*i*_ and *u*_*j*_ are the sinking speeds of the two particles [9]. The rectilinear kernel does not include any distortion of the flow due to the presence of particles, the curvilinear kernel is more accurate and is based on the assumptions of Stokes flow [9]. If we assume our long-distance encounter widths (Fig. 4F) are axisymmetric, our experimental kernels would have the form:

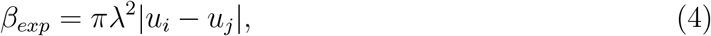

where *λ* is half of the encounter width shown in Fig. 4F. Using this formulation, we find that attached organisms lead to encounter kernels that are always smaller than the rectilinear kernel (Fig. 4H), and ∼ 2× to 10× the curvilinear kernel (Fig. 4I).

### 3.5 *Vorticella* cause sustained rotation of aggregates

Our novel experimental system enabled us to observe aggregates as they sank freely in the ambient fluid without the constraining influence of tethers or other attachments, required in earlier experimental systems [23]. We observed that freely sinking aggregates displayed sustained rotational motion correlated to the presence of *Vorticella* (Fig. 5A, Movie S5). To explore this quantitatively, we developed a custom image-processing pipeline (SI Section 1.2), and used tracked surface fiduciary markers naturally occurring on the aggregates to estimate rotation rates and 3D rotation axes from the 2D images generated by SVTM. We find that aggregates without *Vorticella* show little to no rotation (Fig. 5A). On the other hand, aggregates colonized by one or more *Vorticella* show sustained rotations that are visible by eye even over a 30 s interval (Fig. 5B). To explore if the number of *Vorticella* was the main driver of this rotation, we isolated the effects of aggregate size on the observed rotation rates by rescaling the rotation rate by the scaling factor 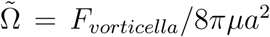, where *F*_*vorticella*_ is the force-scale exerted on the fluid by a single *Vorticella* cell (≈ 200*pN* [26]), *µ* the ambient fluid viscosity, and *a* the aggregate radius. This scaling factor corresponds to the theoretical rotation rate for a spherical aggregate with a single *Vorticella* oriented tangential to the aggregate surface. Upon rescaling, we find that the mean dimensionless rotation rates are higher for aggregates with *Vorticella* than for aggregates without (Fig. 5D; mean dimensionless rotation rate with no *Vorticella*: 0.32 ± 0.27, 1 to 3 *Vorticella*: 0.53 ± 0.27, and greater than 3 *Vorticella*: 0.82 ± 0.50, the differences in mean rotation rates were found to be statistically-significant based on the Kruskal-Wallis test *χ*^2^(3) = 257, *P <* 10^−6^, *N* = 13 distinct aggregates). That the dimensionless rotation rate is 𝒪(1) further indicates that *Vorticella* are indeed the driving factors behind the rotation (Fig. 5D, inset). We further confirmed that measured changes in rotation rates were best explained by *Vorticella* numbers and not due to other experimental conditions (Fig. Supplementary Fig. 2). Interestingly we found that not only does the mean rotation rate rise with *Vorticella* number, but the fluctuations about the mean do as well, with the distribution of higher *Vorticella* numbers becoming heavy-tailed (Fig. 5D and inset).

**Figure 5:**
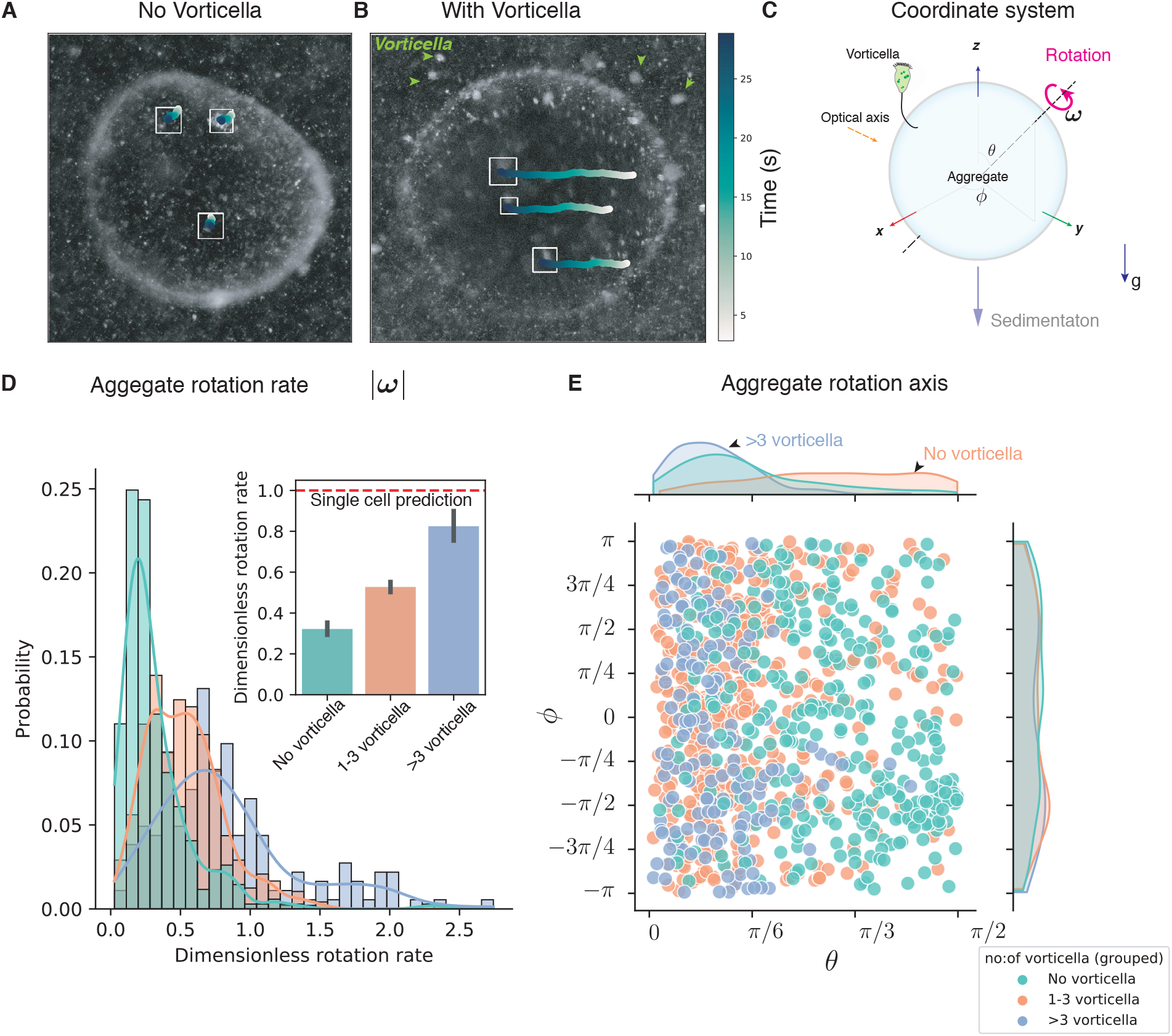
Effect of *Vorticella* on rotational dynamics of sinking aggregates. **(A)** and **(B)** Microscopy snapshots of aggregates without *Vorticella* and with *n >* 7 *Vorticella*, respectively, with tracks of surface features overlaid. Green arrows indicate *Vorticella* locations in (B). Tracked surface features show significant excursion in (B) due to the rotation of the aggregate. **(C)** Coordinate system for quantifying the rotational dynamics of aggregates. Aggregate rotation rate is defined by *ω* and the rotation axis by the polar and azimuthal angle pairs (*θ, ϕ*), respectively. **(D)** Comparison of mean rotation rates of aggregates without and with *Vorticella*. Results are presented over *N* = 7 distinct aggregates for the ‘no *Vorticella*’ condition and over *N* = 6 aggregates for the ‘with *Vorticella*’ conditions. Aggregates with *Vorticella* show significantly larger rotation rates compared to those without *Vorticella* (Kruskal-Wallis test comparing the three conditions *χ*^2^(3) = 257, *P <* 10^−6^). The red-dashed line shows the theoretical rotation rate for an aggregate based on a single *Vorticella*. **(E)** Joint-distribution of the angles that define the axis of rotation. Aggregates without *Vorticella* show a relatively uniform distribution signifying random rotation axis, while aggregates with *Vorticella* tend to rotate about axes that are parallel to the axis of gravity.

We also find that *Vorticella* induce rotations about specific axes that lie close to the axis of gravity, as evident in the distribution of the rotation axis over a unit sphere (Fig. 5C). The distribution is relatively uniform for the no *Vorticella* case, but with increased number of *Vorticella*, the distribution peaks along the axis of gravity (*θ* = 0). We interpret this result as occurring due to gravity breaking the symmetry in orientation space, wherein aggregates have a stable sedimentation orientation along the polar axis (*θ*) due to possible minute density and shape effects that provide a gyrotactic stabilization torque. Thus, the stochastic active torques due to *Vorticella* cause the aggregate to sample orientation space along the azimuthal direction (*ϕ*), as evidenced in our measurements.

## 4 Discussion

Our results suggest a new paradigm for how aquatic sinking aggregates interact hydrodynamically with their surroundings (Fig. 6). In the past, they have been viewed as passive particles, subject to the flow around them, whereas our results show that the flows generated by attached organisms change both aggregate local flow and sinking dynamics. This change is likely to affect every aspect of these particles, including mass transport, the chemical signature they leave in their wake, and the aggregation, dissolution, and settlement rates.

**Figure 6:**
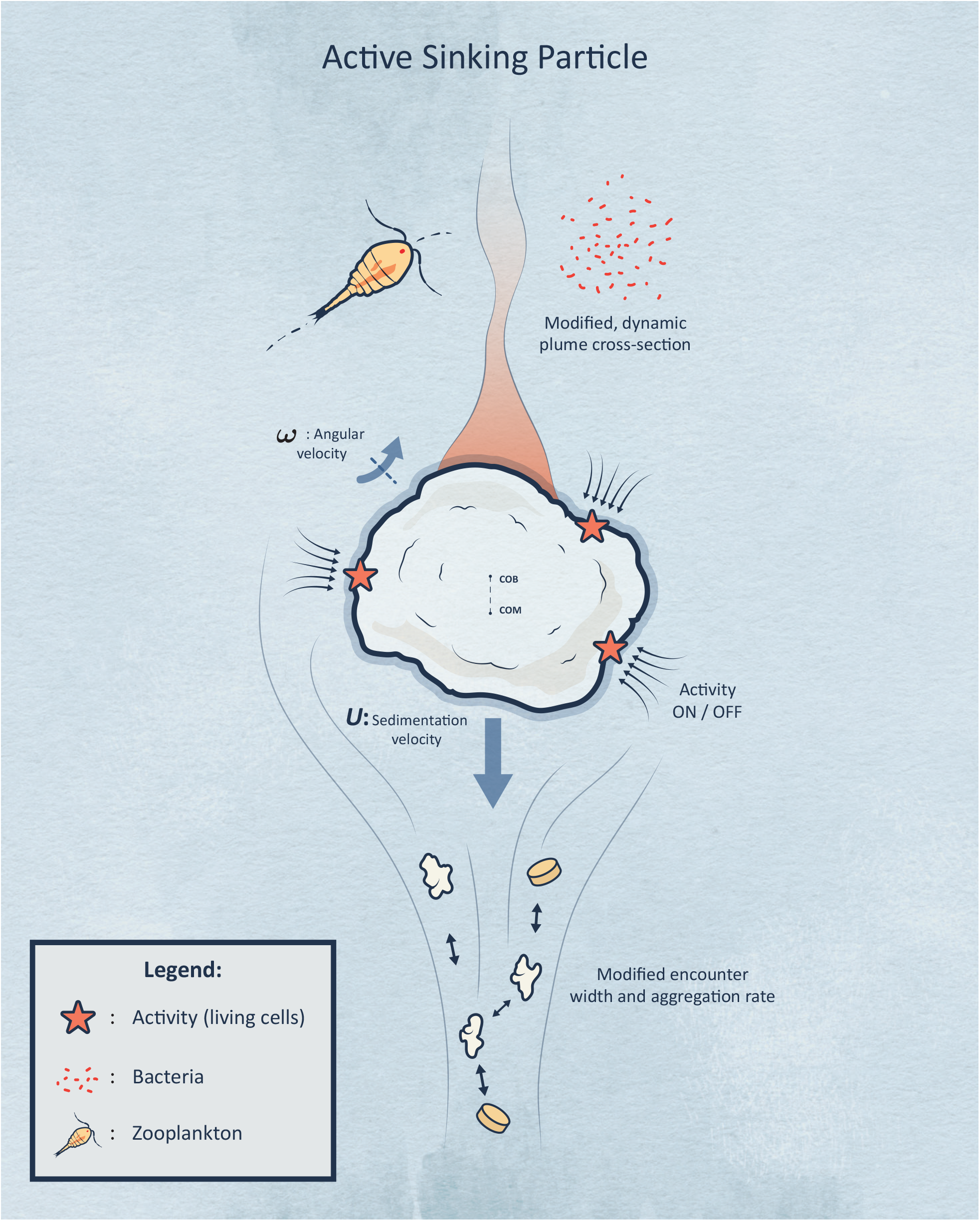
Our results support a new paradigm for “active sinking particles”, compared to the previous view of particles that passively sink with no active hydrodynamic interactions.

Previously, mass transport from small particles or aggregates has been studied in a wide range of flow conditions including uniform flow, shearing flows of different types and, finally, turbulent flow [29, 35, 39, 40, 41]. These works point to the crucial importance of nearfield streamline topology in determining mass transfer (or heat transfer) rates at high Péclet numbers. As shown in this work by direct observation, organisms can change the near-field flow structures significantly, with the flow topology being dependent on the orientation of the activity. This likely leads to a different mass transfer rate scaling than both passive sinking and active swimming particles which scale as Pe^1*/*3^ and Pe^1*/*2^, respectively [29, 41].

The importance of attached organisms’ flow-fields on mass transport was also recently studied in the case of organisms attached to stationary diatoms, wherein the activity induced flows significantly enhanced nutrient fluxes to a simulated diatom cell [34]. Further this enhancement was of a similar scale to that for sinking due to a similar density differential to our particles, hence supporting our finding that even a few attached organisms could change the mass transport characteristics of sinking aggregates. We, thus, anticipate that future investigations of the details of combined advection and diffusion to active sinking particles, and the resulting mass transport rate scaling, may reveal a complex interplay between the two sources of advective transport.

The above changes to mass transport rates would affect how active sinking particles interact with the water column at larger scales. One example of this would be through the plume left behind the particle. Such plumes are an important source of nutrients for microbial life in the water-column and may attract both bacteria [33] and larger organisms like copepods to sinking particles [7, 36, 37]. As shown here, activity significantly modifies the plume cross-section, thus impacting the chances of encounter by bacteria and copepods. In the future, direct visualization of the plume using fluorescent dyes and SVTM in both model and natural aggregates would be an interesting study.

Another example of changes to particles’ interaction with the water column at larger scales is mediated by changes to encounter rates. Studies have shown that changes in near field flows can substantially change encounter rates of sinking aggregates with each other [9, 10]. When combined with coagulation models, moderate changes in encounter dynamics can have large-scale changes on the size spectrum of sinking aggregates as a function of depth [30, 31]. Changes in this size spectrum further significantly change model predictions of carbon sequestration and export to the deep ocean, as well as biogeochemical and trace element distributions in aquatic ecosystems [9, 30, 31]. We find that attached organisms lead to encounter kernels that are always smaller than the rectilinear kernel, and ∼ 2× to 10× the curvilinear kernel. Experimental measurements of particle encounter rates indicate that true coagulation kernels are, indeed, between the rectilinear and curvilinear kernels [42, 43, 9]. This discrepancy has been attributed to porosity, fractal geometry, and intermediate Reynolds number flow [42, 43, 10]. However, our results show that flow from attached organisms could also be a key explanatory factor for this discrepancy. Thus, fully accounting for organism-generated flows my be important for creating accurate models of particle encounters and size spectra in aquatic ecosystems.

Attachment to sinking aggregates also likely affects the feeding rates of attached organisms. Attaching to a sinking aggregate provides organisms with an environment enhanced in food [12, 13, 7, 2, 14] and also changes the hydrodynamics of feeding and organism clearance rates [44, 45]. Previous work with *Vorticella* attached to stationary flat surfaces, showed that ambient flows increase or decrease clearance rates depending on organism orientation relative to flow [26]. While active sinking particles have more complex ambient flow, we similarly observed a diversity of flow speeds and structures near individual *Vorticella*, depending sensitively on the location and orientation of the organisms. This likely leads to clearance rates that are both greater and less than those for organisms on stationary surfaces and that are changing in time as aggregate and organism orientation change.

The activity-induced aggregate rotations discovered in our work mean that the nearfield streamline structures are dynamic in nature. This implies that all of the above effects, including mass transport rates as well as plume and encounter widths, are expected to be dynamic as well. This has interesting parallels to the effects of sub-Kolmogorov scale eddies on particles [40] and sinking aggregates [32], wherein turbulence induced rotations and modifications to the flow around particles can have a substantial effect on mass transport. To understand the relevance of these activity-induced rotations in an ecological setting of the ocean, we can compare the rotation rates from our measurements to those induced by turbulence. The scale of rotation rates seen in our measurements are *F*_*vorticella*_*/*8*πµa*^2^ ≈ 0.03*s*^−1^ (for *a* = 0.5*mm* and *F*_*vorticella*_ = 200*pN*). This corresponds to a velocity gradient due to a turbulence intensity of 10^−9^ *W/Kg*. Compared to turbulence intensities measured in the upper layers of the ocean (away from breaking waves) which range from 10^−10^ to 10^−6^ *W/kg* [46, 47], these activity induced orientation changes may be an important source of orientation stochasticity in the ocean, especially for smaller, more densely colonized aggregates. While outside the scope of this work, our ongoing efforts seek to understand the nature of these dynamics and estimates of the time-averaged mass transport rates, encounter and plume cross-sections. These estimates would then allow connections between lab measurements and ecosystem scale models.

While our results clearly show that attached organisms fundamentally change how sinking aggregates interact hydrodynamically with their environment, our work is just the beginning in understanding both the details of this change and its full implications. Our results are 2D cross-sections of a complex 3D flows; understanding the full 3D flow either through experiment or theory/simulations is an important next step. Indeed, this is possible within the framework of SVTM by leveraging volumetric imaging techniques. Further, we study one size class of sinking aggregate with attached *Vorticella*. Both larger and smaller aggregates are plentiful and important in aquatic ecosystem; similarly, smaller nanoflagellates are abundant on sinking aggregates and generate feeding flows on a smaller scale. Exploring these effects of size scaling of the organisms as well as aggregates is another important area for future investigation. Further, our aggregates, while ideal for a first examination, may be different than real aggregates in several ways. It would be interesting and relevant to investigate how feeding currents of attached organisms interact differently with fractal and irregular aggregates as well as porous aggregates. Our experimental measurements can also serve to validate future flow simulations which can then be used in advection-diffusion models to more fully explore the effects of attached organisms on mass transport to sinking aggregates.

It has recently become increasingly clear that understanding micro-scale processes in aquatic systems is critical for accurately understanding ecosystem level processes [7, 48, 49, 50]. Our results show that it is critical to understand the near-field flow contributions of attached organisms in order to accurately predict the role of sinking aggregates in aquatic ecosystems. By modifying encounter rates, attached organisms may help determine the particle size spectrum of sinking aggregates, and, thus, the many ecosystem processes mediated by aggregates. Similarly, by changing mass transport rates and plume characteristics, attached organisms may modify the composition of bacterial, nanoflagellate, and protist communities living on these aggregates, with follow-on affects on the balance of remineralization and sinking, and, therefore, the export rates of carbon and other nutrients to the deep ocean.

## 5 Materials and Methods

### 5.1 Model sinking aggregates

We created simple model aggregates that share many of the properties of natural aggregates. In nature, macroaggregates are composed of aggregated biological and other debris and range in size from 0.5 mm to several centimeters [1, 5, 2]. Aggregates are typically smaller in estuaries and other areas of high shear (< 2 mm) [2]. In all aquatic environments, aggregates are amorphous and fragile, have varied shapes, and are typically highly porous [1, 5, 6, 2]. Aggregate sinking rates range from less than 1 m day^−1^ to hundreds of m day^−1^, with larger particles sinking faster [5, 2].

We created approximately-spherical particles from 0.3% agarose gel by dripping hot agar into a layer of oil as described in Cronenberg *et al*. [51]. We used vegetable oil rather than kerosene and also made particles of various sizes by directly touching drops to the oil surface while still attached to the syringe needle tip. This density of gel resulted in spheres that sank at velocities within the range observed for similarly-sized marine snow [5]. We next incubated these gel particles overnight in cultures of *V. convallaria*, which were cultured as described in Vacchiano *et al*. [52]. We determined the number of *Vorticella* cells per aggregate by counting them in the microscopy images obtained during Scale-free Vertical Tracking Microscopy.

### 5.2 Flow measurement

We measured flow around active sinking particles using Particle Image Velocimetry (PIV) on short video segments. The water surrounding the spheres was seeded with 2 *µ*m polystyrene spheres (Polysciences 19814). PIV analysis was performed using PIVlab [53]. The aggregates were masked manually; we then used a multi-pass PIV algorithm with decreasing size of the interrogation windows from 128 × 128 pixels to a final window size of 64 × 64 pixels with 50% overlap. Flow fields were then time-averaged over the length of the video (typically, between 1-2 s).

We selected a subset of our particles to analyze that were either bare of *Vorticella*, or that had *Vorticella* in focus and with feeding current primarily directed in the focal plane. We also chose to analyze only video segments where the focal plane coincided with the center of the aggregate in the depth dimension. These choices minimized out of plane flow, so that most of the flows of interest could be measured. As a result of these choices, we analyzed flow fields for 12 video segments across 7 different aggregates (details in SI section Section 1.1). Aggregates ranged in diameter from 0.70 mm to 1.2 mm (mean of 0.97 mm), had sinking speeds of 0.2 mm/s to 5 mm/s (mean of 3 mm/s), and Reynolds numbers of 0.2 to 0.5 (mean of 0.3). Details of each aggregate are in SI Section 1.1, and PIV results for all measured flow fields are available in the Dryad data repository [54].

For obtaining particle pathlines The images were first registered using the Registration plugin in ImageJ [55] to stabilize small movements relative to the camera. The pathlines were then obtained using the FlowTrace plugin for ImageJ [56] by taking the maximum intensity projection of images over a 1 second interval.

### 5.3 Widths of encounter region and plume

Using our measured flow fields, we estimated the width of the plume left behind by the aggregate as well as the width of the encounter region below the aggregate by following streamlines in our measured flow field. This is a reasonable first approach, because the Péclet number (Pe) for these processes is much greater than one, indicating that flow dominates over diffusion in most situations. In the plume, concentrations of small molecules like oxygen and organic solutes, as well as of motile and non-motile bacteria can be increased or decreased by the presence of the aggregate. Diffusion constants for these can range from 10^−9^ m^2^/s (for oxygen and organic solutes) to 10^−14^ m^2^/s (for large non-motile bacteria), with motile bacteria falling in between [6, 28, 57]. Therefore, plume Péclet numbers for our aggregates are ≈ 150 to 10^7^. The Sherwood number, defined as the ratio of total mass transport to mass transport from diffusion alone is also an important indicator of the relative importance of advection and diffusion around a sinking aggregate. For bare sinking spheres of similar Reynolds number to ours, the Sherwood number is 5 for Pe 𝒪 (10^2^) and 10 for Pe 𝒪 (10^4^), again indicating that advection dominates over diffusion [6].

When considering the encounter region below the sphere, advection is even more dominant, and the Péclet number even higher. For instance, considering the encounter rate with a 100 *µ*m sphere, the Stokes-Einstein relation gives D = 2 × 10^−15^ m^2^/s with a resulting Péclet number of 10^8^ [58].

When finding the plume width, we started streamlines in a ring around the aggregate at a distance of 140 *µ*m from the aggregate surface and integrated forward in time in the measured flow field (Fig. 4). Streamlines that approached within 16 pixels (35 *µ*m) of the aggregate surface were terminated, as the flow field measurement is noisy in this region. The width of the plume at each height above the aggregate is the distance between the outer-most streamlines. Here, choosing a distance of 140 *µ*m from the aggregate surface can be considered choosing an approximate diffusion constant for the substance of interest: from a scaling perspective, the substance will diffuse approximately 140 *µ*m in the time it takes the aggregate to sink its own diameter. This yields *D* ∼ 10^−9^ m^2^/s, appropriate for oxygen or small organic solutes.

Similarly, we estimated the width of the encounter region below the aggregate, following the ideas in Humphries *et al*. [10]. Particles within this region of fluid would come in to contact with the aggregate as it falls through the water column. Similar to our plume calculation, we determined streamlines that began in a ring around the aggregate at a distance of 140 *µ*m from the aggregate surface. To find the encounter region, we then integrated backward in time in the measured flow field. The width of the encounter region is the distance between outer-most streamlines (equivalent to 2 × *λ* in Humphries *et al*. Fig. 2 [10]). Choosing a beginning location of 140 *µ*m from the aggregate surface effectively gives the encounter region for particles in the water column that are 140 *µ*m or larger.

We compared our measured plume and encounter region widths to the known results for Stokes flow and Oseen’s modification to Stokes flow ([38] eqns. 4.9.12 and 4.10.3), where the Oseen flow was calculated for Reynolds numbers that matched parameters for each individual aggregate (see SI Table 1).

For more direct comparison with work such as Humphries *et al*. and with theoretical coagulation kernels [9, 10], we extended our experimentally measured streamlines beyond the field of view of our experiments to a final vertical distance above and below the sphere of 20 times the particle radius (20*a*) using Oseen flow around a sinking sphere. We began the theoretical streamlines at the location of our experimentally-measured streamlines when they were a vertical distance of 2*a* above or below the middle of the sphere (above for plume measurements and below for encounter region). The width of the plume at 20*a* was, then, the distance between the outer-most theoretical streamlines. These long-distance widths were only calculated for particles where the distance 2*a* was in the field of view (N=9 for encounter volume and N=10 for plume).

## Authors’ Contributions

D.K., M.P, and R.E.P. designed research; D.K and R.E.P. carried out experiments; D.K. and R.E.P. carried out the analysis. D.K. and R.E.P. wrote the paper with comments from M.P.

## Data Availability

Original PIV flow field data for all included flow fields are available in the Dryad data repository [54].

## 6 Acknowledgements

We thank Rebecca Konte for graphics and artwork in Fig.1 and 6. We are grateful to A. Andersen, E. Riley, and T. Kiørboe for helpful discussion. We also thank Rahul Chajwa, Hongquan Li and Prakash Lab members for valuable discussions. D.K acknowledges support from Bio-X fellowship and Schmidt Science Fellowship. We thank Hopkins Marine Station for lab-space for experiments. We are also grateful for support from NSF grant # IOS-1755326 to REP. M.P. acknowledges financial support from NSF Career Award, Moore Foundation, HHMI Faculty Fellows program, NSF CCC (DBI-1548297) program, NSF Convergence Award (OCE-2049386), Schmidt Foundation and CZ BioHub Investigators program.

