## Supplemental Information for "Active Sinking Particles: Sessile Suspension Feeders significantly alter the Flow and Transport to Sinking Aggregates"

### Supplementary Information

Deepak Krishnamurthy [*,a,1], Rachel Pepper [*,b,1], Manu Prakash[a]

[a]Department of Bioengineering, Stanford University, Stanford, California, USA

[b]Department of Physics, University of Puget Sound, Tacoma, Washington, USA

[*] Equal contribution

[1]To whom correspondence should be addressed;

August 5, 2021

**3 Supplementary Movie Captions** 5

| Figure panel | Particle ID | Number of *Vorticella* | a (mm) | U (mm/s) | Re |
|---|---|---|---|---|---|
| **-** | **1** | **0** | **0.38** | **0.34** | **0.3** |
| A | 2 | 0 | 0.52 | 0.46 | 0.5 |
| B | 3 | 1 | 0.35 | 0.22 | 0.2 |
| C | 3 | 1 | 0.38 | 0.21 | 0.2 |
| **-** | **4** | **3** | **0.49** | **0.20** | **0.2** |
| D | 4 | 3 | 0.49 | 0.21 | 0.2 |
| E | 4 | 3 | 0.49 | 0.22 | 0.2 |
| **-** | **5** | **3** | **0.49** | **0.21** | **0.2** |
| F | 5 | 3 | 0.49 | 0.23 | 0.2 |
| G | 6 | 3 | 0.57 | 0.45 | 0.5 |
| H | 7 | 7 | 0.59 | 0.37 | 0.4 |
| I | 7 | 7 | 0.61 | 0.41 | 0.5 |

**Supplementary Table 1** Parameters for all measured flow fields the "Figure panel" column refers to the panel where the flow field is shown in Supplementary Fig. 1. Those with a dash in this column are shown in the main text Fig. 3

### 1 Supplementary Methods

#### 1.1 Experimentally-Measured flow fields

We measured flow at 12 different time point around seven sinking aggregates. These 12 flow fields were used to determine plume and encounter widths (Sec . 5.3, Main text). Aggregate parameters are listed in Table 1 and flow measurements are in Supplementary Fig. 1. Full PIV results for all measured flow fields are available in the Dryad data repository [1].

#### 1.2 Computing an object's rotation rate and axis from 2D images

To quantify the rotation rates of the aggregates, we solved the problem of how to estimate the 3D rotation vector from the 2D images of the aggregates/spheres. Our idea is based on using tracked surface features to estimate the rotation rate and axis by using the fact that the aggregate shape is approximately spherical. Surface features naturally occurring on the aggregate surface served as excellent fiduciary markers and we tracked these features over time using a custom image-processing pipeline. Briefly, we selected at least 3 surface features manually and then tracked them automatically over subsequent frames using the OpenCV CSRT tracker ([2]), implemented in a custom Python pipeline. We also concurrently tracked the whole aggregate (using the OpenCV KCF tracker) in order to get an accurate localization of the aggregate centroid. Tracks of the surface features and aggregate centroid were then used to estimate the rotation rate and axis using a least-squares method. We briefly describe the method below:

Velocity of any point on a sphere rotating about an arbitrary axis at a rate $\omega$ is given by $\boldsymbol{v} = \boldsymbol{\omega} \times \boldsymbol{r}$. In a cartesian coordinate system, we can take the projection of this on the XZ plane (image plane), which gives:

$$v_x\hat{x} + v_z\hat{z} = (\omega_y r_z - \omega_z r_y)\hat{x} + (\omega_x r_y - \omega_y r_x)\hat{z} \tag{1}$$

We have 3 unknowns $(\omega_x, \omega_y, \omega_z)$ (or equivalently, the rotation rate $|\omega|$ and the axis), so we need at least 3 equations to have a solution. However, the above only gives us two equations (for x and z). We can make progress by tracking more features on the sphere simultaneously. In general, we can write the above as a linear system for $M$ tracked points as:

$$\begin{bmatrix} 0 & r_{1,z} & -\sqrt{R^2 - r_{1,x}^2 - r_{1,z}^2} \\ \sqrt{R^2 - r_{1,x}^2 - r_{1,z}^2} & -r_{1,x} & 0 \\ 0 & r_{2,z} & -\sqrt{R^2 - r_{2,x}^2 - r_{2,z}^2} \\ \sqrt{R^2 - r_{2,x}^2 - r_{2,z}^2} & -r_{2,x} & 0 \\ \vdots & \vdots & \vdots \\ 0 & r_{M,z} & -\sqrt{R^2 - r_{M,x}^2 - r_{M,z}^2} \\ \sqrt{R^2 - r_{M,x}^2 - r_{M,z}^2} & -r_{M,x} & 0 \end{bmatrix} \begin{bmatrix} \omega_x \\ \omega_y \\ \omega_z \end{bmatrix} = \begin{bmatrix} v_{1,x} \\ v_{1,z} \\ \vdots \\ v_{M,x} \\ v_{M,z} \end{bmatrix} \tag{2}$$

Note that by assuming the shape of the object, we can estimate the out-of-plane location which is necessary to define the linear operator on the left. We also assume that we are on one particular side of the sphere (either towards or away from us). Since the sphere can be transparent our estimate for the angular velocity vector will have an uncertainty in sign, except when the rotation is mostly along the optical axis. From the above equation we have $2M$ equations and 3 unknowns, which we solve as a least-squares minimization problem.

We validated the method by generating test-data of aggregates with arbitrary rotation rates and orientations and comparing the values estimated by our method to the ground-truth (Supplementary Fig. 3). As seen in Supplementary Fig. 3 the method performs effectively and accurately estimates both rotation rates and axis for diverse cases including fully 3D rotation vectors, where the axis does not lie along any principal axes in relation to the camera axis. Over all the orientations we considered the relative errors in rotation rate estimates were $< 1\%$, and relative errors in the rotation axis were $<= 2\%$. This serves as validation for this method. This method was then used to estimate the rotation rates and axis from our Scale-free Vertical Tracking Microscopy data of sinking aggregates (Fig. 5, Main text).

### 2 Supplementary Discussion

#### 2.1 Quantifying the effects of walls and other experimental parameters on the dynamics of aggregates

To confirm if *Vorticella* were indeed the main cause of the observed increase in rotation rates of the freely sinking aggregates, we systematically measured the effects of other experimental variables. We compared the dimensionless rotation rates of the aggregates with respect to distance to radial walls (Supplementary Fig. 2A), distance to axial walls (Supplementary Fig. 2B), aggregate size (Supplementary Fig. 2C), and found no significant correlations ($R^2$ values for these cases was $< 0.004$). In striking contrast, a clear correlation is seen when looking at the dimensionless rotation rate as a function of no:of vorticella (Supplementary Fig. 2D) ($R^2 = 0.32$). This confirms that the observed rotational dynamics is indeed due to the *Vorticella* .

#### 3 Supplementary Movie Captions

**Supplementary Movie 1** Multi-scale measurements of freely-sinking aggregates with *Vorticella* (Active Sinking Particles) using Scale-free Vertical Tracking Microscopy and model aggregates.

**Supplementary Movie 2** Flow around a freely sinking aggregate with no *Vorticella* .

**Supplementary Movie 3** Flow around a freely sinking aggregate with 1-3 *Vorticella* .

**Supplementary Movie 4** Flow around a freely sinking aggregate with $> 3$ *Vorticella* .

**Supplementary Movie 5** Activity of *Vorticella* induces rotation of the aggregates as they sink.

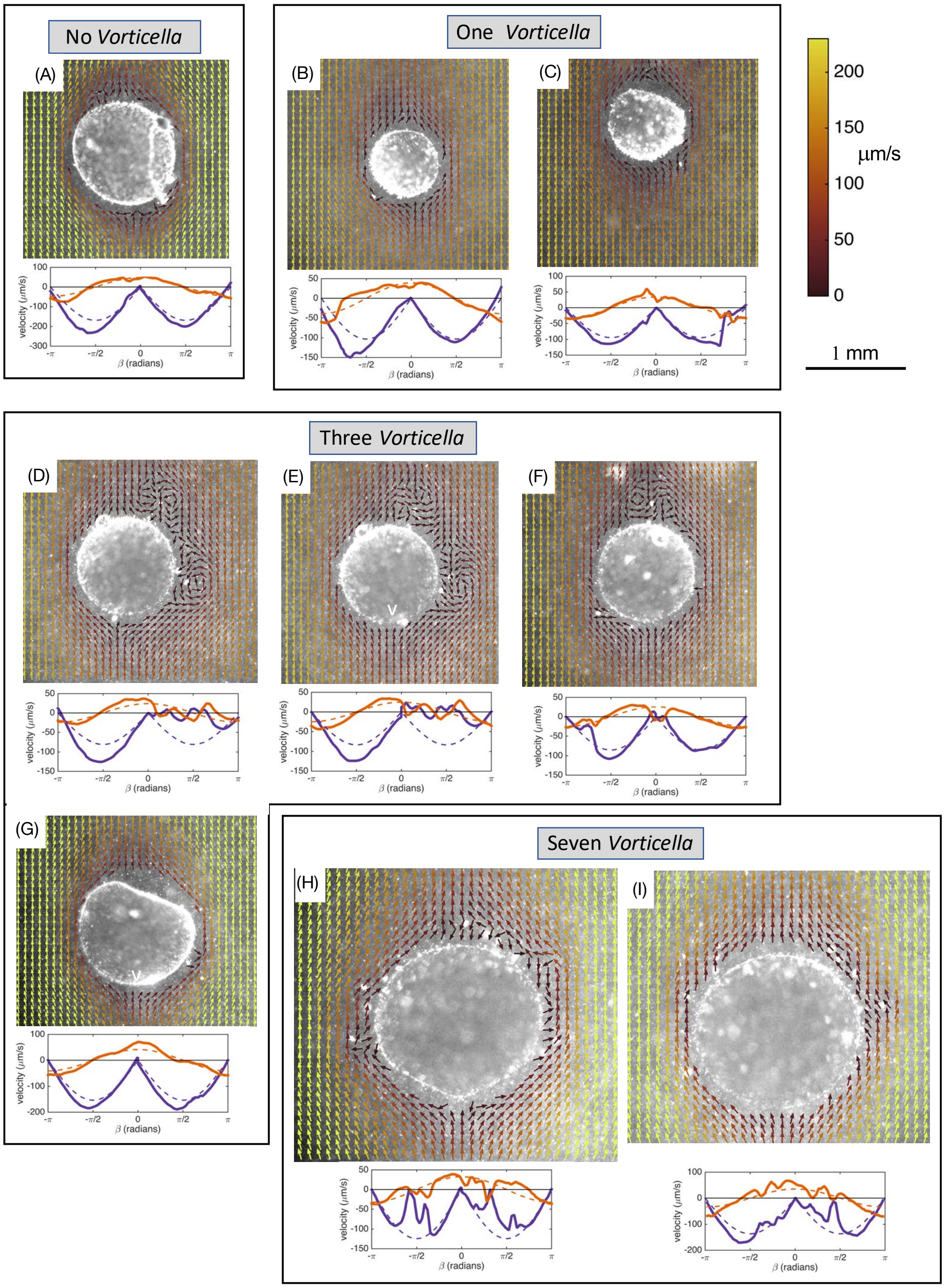

**Supplementary Fig. 1**

**Supplementary Figure 1**: **Flow around active sinking aggregates. Upper image** for each panel shows measured flow velocities. Arrows indicated flow direction and color indicates speed. The scales on the far right apply to all panels. **Lower image** shows measured velocities (solid lines) along a circle 200 $\mu$m from the sphere surface (e.g. blue line in panel main text Fig. 3(B)). $\beta$ is the polar angle indicated in panel main text Fig. 3(B), and is positive to the right of the sphere and negative to the left. Perpendicular and parallel components of the velocity are relative to the sphere surface. Dashed lines are calculated velocities for Stokes flow (zero Reynolds number) around a sphere. Relevant parameters for each aggregate are listed in Table 1.

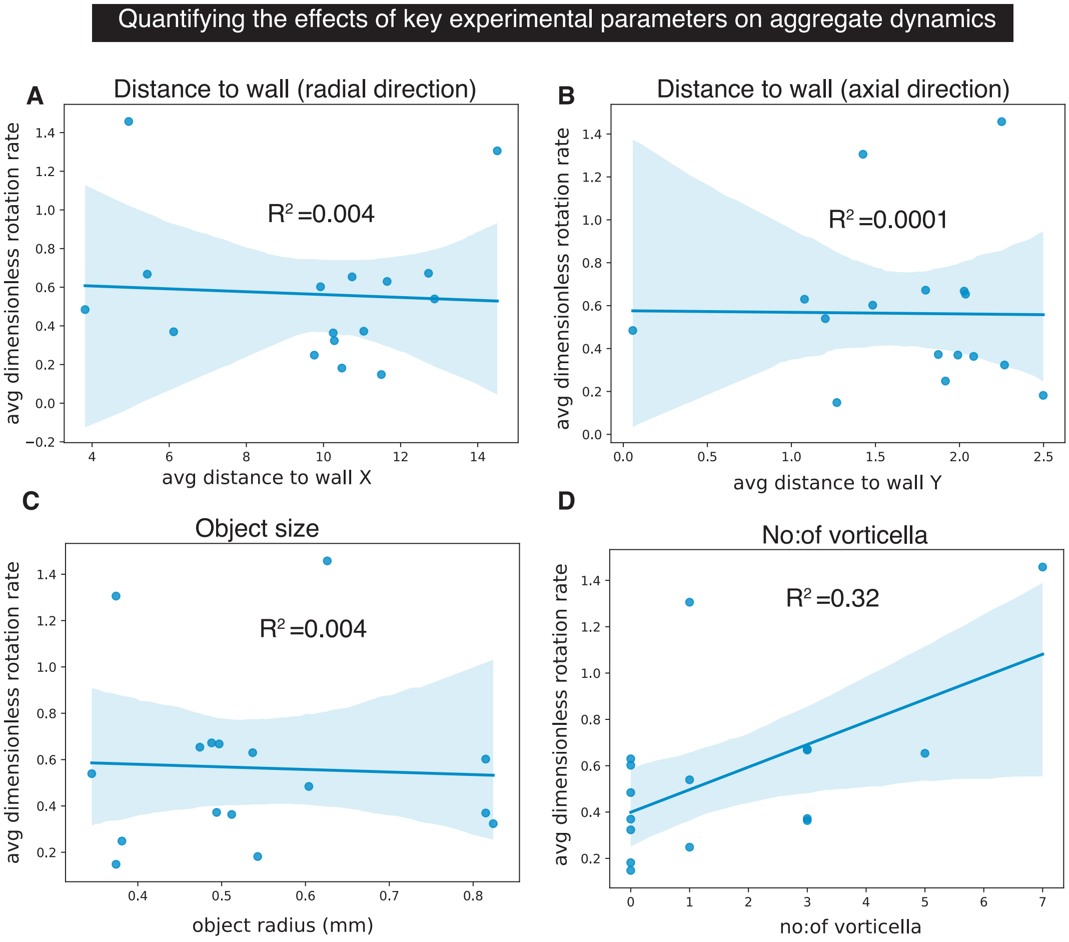

**Supplementary Fig. 2**

**Supplementary Figure 2**: **Quantifying effects of key experimental parameters on the measured rotational dynamics of the aggregate**. Dimensionless rotation rate of aggregates averaged over a track and plotted with respect to **(A)** Distance from radial walls,**(B)** distance from axial walls, **(C)** the aggregate size and **(D)** the no:of vorticella attached to the aggregate. The dots represent averages over each measured trajectory, the solid line represents the least-squares linear fit and the shaded region the 95% confidence interval of the fit.

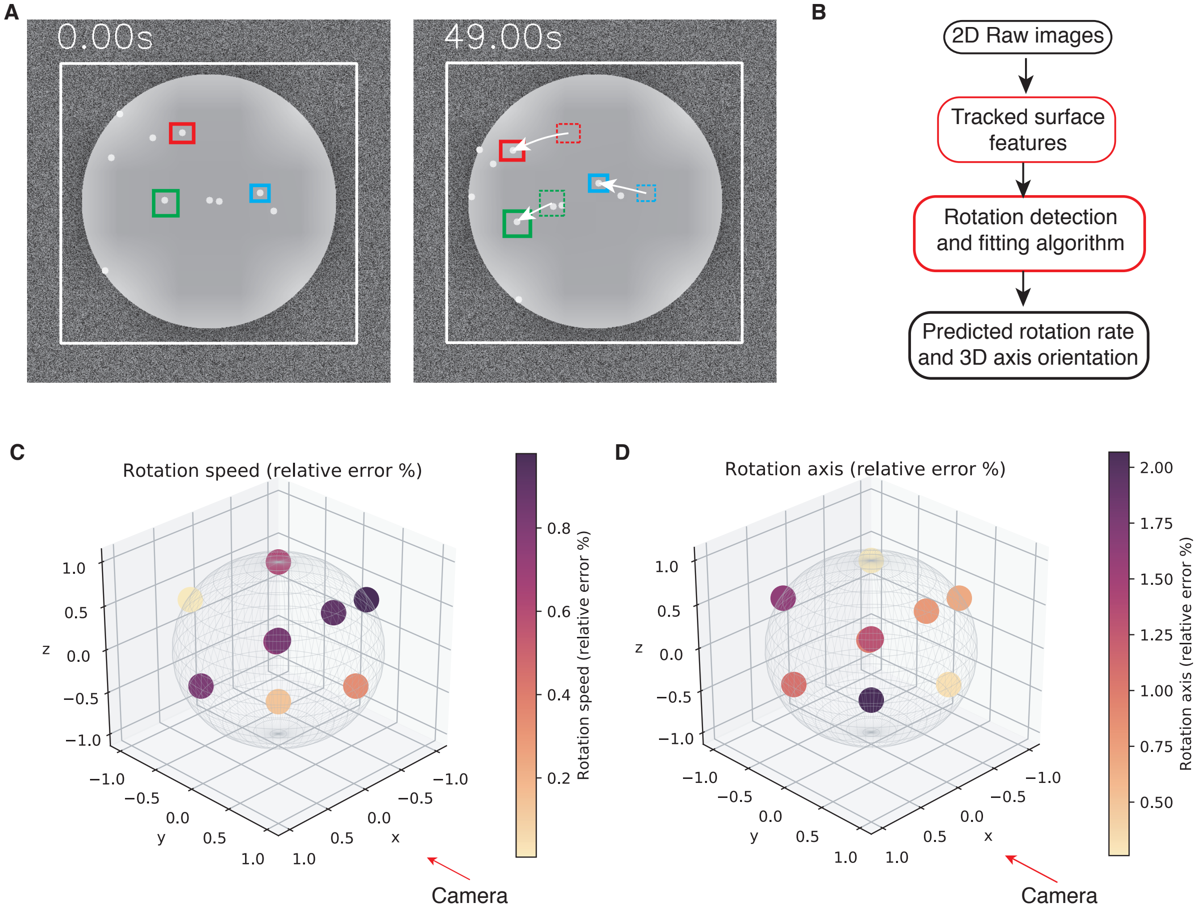

**Supplementary Fig. 3**

**Supplementary Figure 3**: **Validation of the rotation analysis**. **(A)** We used computer-generated test-images of aggregates with surface features to simulate object rotations about arbitrary 3D axes in relation to the camera. Randomly distributed features on the object surface (white dots) served as fiduciary markers and were tracked over time (colored boxes). **(B)** The images were run through the rotational analysis pipeline to predict both the solid-body rotation rate and 3D rotation axis. **(C)** and **(D)** Relative error between actual and predicted rotation speeds and rotation axes for a wide range of rotation vectors. The camera is looking along the Y-axis and the image plane is the XZ plane.
